# The pecten oculi regulates pH in the bird eye

**DOI:** 10.64898/2026.11.24.768906

**Authors:** Anna Victoria G. T. Mikkelsen, Morten Hulbæk Fog, Amanda Bundgaard, Freja Terndrup Kristensen, Thomas Vorup-Jensen, Tobias Wang, Catherine J.A. Williams, Christian Damsgaard

## Abstract

The bird retina generates a dual acid load: aerobic metabolism in the outer retina produces CO_2_, while chronic anoxia in the inner retina yields lactic acid, raising the question of how intraocular pH homeostasis is maintained in birds. We show that chicken (*Gallus gallus*) vitreous pH remains close to blood pH despite these respiratory and metabolic acid-base challenges and a low non-bicarbonate buffering capacity, which is compensated by high vitreous bicarbonate. Spatial transcriptomics of eye tissue revealed enriched expression of carbonic anhydrase 2 and 4, and vacuolar-type H^+^-ATPase within the intraocular vascular structure, the pecten oculi. Immunohistochemistry confirmed the presence of carbonic anhydrase 2 in the pecten, suggesting an important role in ocular CO_2_ removal and pH regulation. Together, our findings demonstrate that bicarbonate buffering and pecten-mediated exchange of acid-base-relevant ions and metabolites act in concert to sustain retinal function under the extreme metabolic conditions of the bird eye.

**Summary statement:** The avascular bird retina produces both CO_2_ and lactic acid, yet the vitreous pH remains close to blood pH via pH regulation in the pecten oculi.

## Introduction

Few tissues rival the metabolic demands of the vertebrate retina, which consumes oxygen and glucose at mass-specific rates exceeding even those of the brain (Ames III, 1992; Country, 2017; Damsgaard and Country, 2022). In mammals, this demand is met through a dual blood supply with choroidal vessels perfusing the outer retina, while intraretinal capillaries deliver oxygen directly to the inner layers (Buttery et al., 1991; Nickla and Wallman, 2010). This vascular arrangement, however, imposes an inherent opto-respiratory compromise, as blood vessels and nerve fibres positioned anterior to the photoreceptors scatter incoming light, reducing optical clarity (Damsgaard and Country, 2022). Retinal thickness and vascularisation are thus constrained by competing demands for metabolic support and optical transparency.

The bird retina represents an extreme case of this compromise, as it entirely lacks intraretinal blood vessels, yet reaches thicknesses of 400–600 µm, far exceeding the ∼150 µm oxygen diffusion limit from the choroid (Damsgaard and Country, 2022; Dollery et al., 1969; Potier et al., 2020). This anatomical arrangement supports high cell densities for light capturing and visual processing, but raises the question of how this metabolically active tissue achieves oxygen and nutrient supply as well as waste product clearance without internal blood perfusion. Recent data document continuous anoxia and glycolytic ATP production in the inner retina (Damsgaard et al., 2026). This metabolic phenotype generates a substantial acid load, where the oxygenated outer retina continuously produces CO_2_ and the anoxic inner retina produces lactic acid. Both acidifiers accumulate in the vitreous humour (Damsgaard et al., 2026; Marcinek et al., 2010; Robergs et al., 2004). In the absence of intraretinal blood vessels, neither CO_2_ nor lactic acid can be cleared by a local circulation, raising the risk of retinal and ocular acidification.

One of the key candidates for managing the acid load is the pecten oculi, a highly vascularised intraocular structure unique to birds (Brach, 1977; Meyer, 1977). It is proposed to facilitate exchange of glucose, CO_2_, and lactate between the retina and the blood via the vitreous humour (Damsgaard et al., 2026). A pecten ablation study in chicken suggested a role of the pecten in pH regulation (Brach, 1975), but the mechanisms underlying pH regulation and the physicochemical buffering capacity of the vitreous are not known. We therefore aimed to identify the mechanisms underlying pH regulation in the bird eye. Specifically, we examined the *in vivo* and *ex vivo* acid-base properties of the vitreous humour and the expression of pH-relevant transporters and enzymes in the pecten oculi, providing integrative insight into how pH is regulated under the extreme respiratory conditions of the bird eye.

## Materials and methods

### Animal husbandry

Commercial chickens (*Gallus gallus domesticus,* Dekalb White) were obtained as 17-week-old pullets from a Danish commercial producer. These birds originated from EU-based commercial rearing, ensuring typical early-life conditions and genetic background. Upon arrival at Aarhus University, Research Centre Foulum (Viborg, Denmark), hens were housed at a density of 4 hens m^−2^ in a multitier aviary system (4 x 3.2 x 2 m) equipped with nest boxes, perches and floor substrate, with ad libitum access to water and standard commercial pre-lay and subsequent layer feed (DLG), with supplemental calcium shells. Temperature and light programmes (artificial warm white) were in line with commercial recommendations. Animals were monitored twice daily by trained personnel. All procedures involving this cohort were approved by the Danish Animal Experiments Inspectorate (permit no. 2024-15-0201-01744).

For immunohistochemistry, fertilised eggs of chickens (Dekalb White) were obtained from Topæg (Viborg, Denmark) and transported to the Department of Biology, Aarhus University. Eggs were incubated in a 24-egg digital incubator (cat.no. 738; America A/S, Thisted, Denmark) following the manufacturer’s protocol, and newly hatched chicks were subsequently transferred to a h x l x w = 2 x 2 x 1 m^3^ indoor aviary maintained at 22°C with continuous access to food and water. Experimental procedures on these birds were approved by the Danish Animal Experiments Inspectorate (permit no. 2022-15-0201-01278).

### *In vivo* acid-base status

Plasma and vitreous samples were collected from eight 45-week-old female laying hens (body mass 1.75 kg, s.d. = 0.14) at AU Viborg, Research Centre Foulum.

Blood was sampled from the right jugular vein of a non-anaesthetised, restrained chicken and collected in EDTA tubes (BD Vacutainer, Becton Dickinson, Franklin Lakes, NJ, USA) to prevent clotting. Vitreous humour samples were collected after euthanasia by an intravenous overdose of sodium pentobarbital (400 mg mL^−1^, 0.7–1 mL per hen; Exagon Vet, Richter Pharma, Wels, Austria) administered via the left jugular vein. The vitreous humour from both eyes of each hen was pooled. Immediately after sampling, pH was measured in blood and vitreous humour at 42°C with a pH meter (SevenCompact™ S210, Mettler Toledo, Greifensee, Switzerland). After the pH measurements, blood was centrifuged (10 min, 2122 g, 4°C) to isolate plasma, and plasma and vitreous humour were stored at −70°C until further analysis. EDTA tubes were used because the samples were also used for other analyses; in paired heparinised and EDTA plasma samples (n = 4), EDTA had no significant effect on [Na^+^] (*P* = 0.60), [Cl^−^] (*P* = 0.10) or total CO_2_ (*P* = 0.50; paired t-tests). As EDTA may slightly lower measured blood pH, blood pH values may be marginally underestimated.

### Vitreous acid-base properties

#### Tonometry

To determine the non-bicarbonate buffer capacity and the apparent pK of carbonic acid experimentally, plasma and vitreous samples from four hens from the same cohort (body mass 1.67 kg, s.d. = 0.07) were gas-equilibrated in an Eschweiler rotatory tonometer system within a water bath maintained at 42°C. The tonometers received humidified gas mixtures from a gas-mixing pump (Type 2M301/a-p; H. Wösthoff GmbH, Bochum, Germany), allowing precise adjustment of the partial pressures of N_2_, O_2_, and CO_2_ (pN_2_, pO_2_, and pCO_2_, respectively). Gas compositions were 20% O_2_ combined with 2% (14 mmHg), 4% (28 mmHg), or 8% (56 mmHg) CO_2_, with N_2_ making up the balance. Samples were allowed to equilibrate for 30 min between steps. For pH determination, 0.5 mL of the equilibrated sample was withdrawn using a syringe and measured immediately with a temperature-controlled pH meter (Mettler Toledo). Additionally, a portion of the equilibrated sample was collected using a custom-made Hamilton syringe to determine total CO_2_ concentration ([CO_2_]_total_) (see details below) and [HCO_3_^−^] was calculated as:

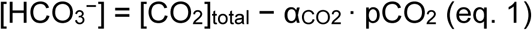

where [CO_2_]_total_ represents the sum of dissolved CO_2_ and bicarbonate, α_CO2_ represents the solubility coefficient of CO_2_ (0.028 mmol L^−1^ mmHg^−1^ at 42°C) (Chiodi and Terman, 1965), and pCO_2_ is the water-vapour-corrected chamber pCO_2_ for each equilibration step. Based on these data, the non-bicarbonate buffer capacity (β_NB_) was calculated as:

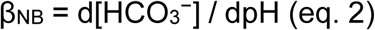

The apparent dissociation constant (pK’) of carbonic acid was calculated for each measurement as:

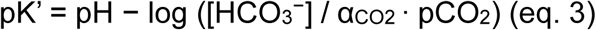

#### Ion, protein, and metabolite composition

We next compared the ion, protein and metabolite composition of the plasma and vitreous humour, which were thawed on ice before analysis (n = 8). Bicarbonate concentration was measured using a custom setup for CO_2_ quantification based on Hamad et al., 2024 and Lee et al., 2018. The setup consisted of a self-designed, resealable funnel containing 1.5 mL of 0.01 mol L^−1^ HCl, which was continuously bubbled from the bottom with N_2_ gas at a constant flow rate of 200 mL min^−1^. The sample was injected into the funnel, where the H^+^ from HCl converted all HCO_3_^−^ into CO_2_ gas.

The released CO_2_ was carried through a condenser on ice and CaCl_2_ beads to remove water vapour, and the CO_2_ content was quantified by infrared detection using an LI-7000 CO_2_/H_2_O analyser (LI-COR Biosciences, Lincoln, NE, USA). The total CO_2_ content was quantified by integrating the area under each CO_2_ response curve. After manual baseline correction of the raw data, the area under the curve was calculated for each predefined injection interval, including the 10 mM and 30 mM NaHCO_3_ standards as well as the vitreous and plasma samples.

The experimentally derived pK’ from the tonometer experiments was combined with the measured *in vivo* pH and total CO_2_ to calculate *in vivo* pCO_2_ using the rearranged Henderson–Hasselbalch equation:

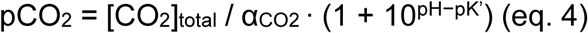

The *in vivo* [HCO_3_^−^] was then calculated with eq. 1 from the *in vivo* pCO_2_ obtained with eq. 4.

To estimate what blood pH would have been under the vitreous acid load, we calculated for each hen the blood pH at the vitreous pCO_2_ and lactate concentration using the Henderson–Hasselbalch equation:

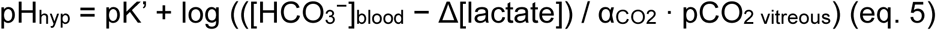

where Δ[lactate] is the excess vitreous lactate ([lactate]_vitreous_ − [lactate]_plasma_), assumed to titrate HCO_3_^−^ 1:1. In hens in which Δ[lactate] exceeded blood [HCO_3_^−^], bicarbonate would be fully titrated and pH_hyp_ could not be calculated.

Sodium concentration was measured using a flame photometer (Sherwood Scientific, Cambridge, UK). Chloride concentrations were determined using an MKII Chloride Analyzer (Sherwood Scientific). Total protein concentrations were quantified using the Pierce™ 660 nm Protein Assay (Thermo Fisher Scientific, Waltham, MA, USA) in triplicate. Phosphate and lactate concentrations were measured using commercially available colourimetric assay kits (MAK488 and MAK329, respectively; Sigma-Aldrich/Merck, St. Louis, MO, USA) according to the manufacturer’s instructions.

### Spatial transcriptomics analysis

To identify candidate genes linked to pH regulation, we re-analysed a published spatial transcriptomics data set from one chicken eye, consisting of raw and aligned spatial RNA-seq data available at the Gene Expression Omnibus (GEO) (GSE309408) (Damsgaard et al., 2026). Analyses were conducted in R (R Core Team, 2026) using the Seurat package (Hao et al., 2021). To ensure the data were comparable across different spots and samples, normalisation was performed using the *NormalizeData*()-function. The analysis focused on genes involved in pH regulation, including those encoding proton transporters, bicarbonate transporters, associated enzymes, or regulatory proteins. Candidate genes were identified through a review of scientific literature and supplemented by queries to the GeneCards, NCBI Gene, and Ensembl databases. Differential gene expression between the pecten and the other ocular tissues was tested using a Wilcoxon rank-sum test with the FindMarkers() function in Seurat and reported as log_2_ fold change (log_2_FC) and −log_10_(adjusted P), with Bonferroni correction across all genes.

### Immunohistochemical validation

To validate the spatial transcriptomics findings at the protein level, immunohistochemistry was performed on selected highly expressed gene products. As no antibodies raised against the chicken proteins were commercially available, we used antibodies raised against mammalian proteins (sheep polyclonal anti-CA2, PA5-48006; rabbit polyclonal anti-CA4, 13931-1-AP; mouse monoclonal anti-ATP6V1C2, clone OTI3E9, MA5-26964; all Invitrogen, Thermo Fisher Scientific, Carlsbad, CA, USA), which are validated in mammals but not in birds. Their specificity in chicken tissue was therefore tested by Western blotting. The CA2 antibody gave a single band at the expected molecular weight (Fig. S1), whereas the antibodies against CA4 and ATP6V1C2 did not give a specific band. Immunohistochemical data are therefore reported for CA2 only.

#### Tissue preparation

Three chickens (one female and two males; body mass 0.19 kg, s.d. = 0.011) aged 7–9 days were anaesthetised using isoflurane (5% in air) delivered at a flow rate of 3 L min^−1^ via a vapouriser (E-vet, Langeskov, Denmark). Deep anaesthesia was maintained for 3 to 4 min, after which animals were euthanised by decapitation. Following euthanasia, the eyelid was gently opened, and a small orientation mark was made at the top of the eye using a heated tip, and the eye was removed. Each eye was placed in a polyethylene 19×13 mm embedding mould (TAAB, Aldermaston, UK) filled with optimal cutting temperature compound (Tissue-Tek, Torrance, CA, USA). Eyes were oriented for transverse midline sectioning. Moulds were snap-frozen by immersion in isopentane pre-cooled in liquid nitrogen for 1–3 min and stored at −70°C. Cryosectioning was performed using a cryostat (Leica CM1950, Leica Biosystems, Deer Park, IL, USA) with both block and knife temperature at −20°C, providing 10 µm thick transverse sections near the pecten. Tissue sections were carefully transferred to Superfrost™ Plus microscope slides (REF: J1800AMNZ, Thermo Fisher Scientific) and stored at −70°C.

#### Immunohistological staining and imaging

Tissue sections were equilibrated to room temperature and fixed by immersing the slides in 3.7% (w/v) formaldehyde (diluted 1:10 from a 37% formaldehyde solution; Cat# F8775, Sigma-Aldrich/Merck) in phosphate-buffered saline (PBS, pH 7.4; Cat# P4417, Sigma-Aldrich/Merck) for 10-15 min, followed by three 5 min washes in PBS. Permeabilisation and blocking were then performed by incubating the sections for 1 h in PBS containing 0.3% (v/v) Triton X-100 (Cat# STBJ3368, Sigma-Aldrich/Merck), 2% (v/v) chicken serum (Cat# C5405, Sigma-Aldrich/Merck), 10% (v/v) donkey serum (Cat# 0030-01, SouthernBiotech, Birmingham, AL, USA) and 3% (w/v) bovine serum albumin (BSA; Cat# A7030-10G, Sigma-Aldrich/Merck). After blocking, slides were again washed three times in PBS for 5 min each. The antibody diluent consisted of PBS with 1% (w/v) BSA, 0.05% (v/v) Tween-20 (Cat# P1379, Sigma-Aldrich/Merck) and 0.1% (v/v) donkey serum. The primary antibody, polyclonal sheep anti-CA2 (Cat# PA5-48006, RRID: AB_2609833; Invitrogen, Thermo Fisher Scientific), was diluted 1:41 in antibody diluent to a final concentration of 4.88 µg mL^−1^. Sections were thereafter incubated overnight at 4°C with antibody diluent containing the primary antibody. Following overnight incubation, sections were washed three times in PBS for 5 min. Secondary antibody incubation was then performed for 1 h at room temperature in the dark with polyclonal donkey anti-sheep IgG conjugated to Alexa Fluor 647 (Cat# A-21448, RRID: AB_2535865; Invitrogen, Thermo Fisher Scientific) diluted 1:1000 in antibody diluent to a final concentration of 2.00 µg mL^−1^. Sections were then washed three times in PBS for 5 min each, then mounted with DAPI Fluoromount-G® (Cat# 0100-20, SouthernBiotech) and a coverslip, and sealed with clear nail polish before imaging. Controls included a DAPI-only negative control and a secondary antibody-only control, in which the secondary antibody was applied without primary antibody to test for non-specific binding. Experimental samples were stained with both DAPI and anti-CA2.

Immunofluorescence images were captured on a Zeiss LSM 900 laser scanning confocal microscope equipped with an Airyscan 2 detector (Carl Zeiss Microscopy, Jena, Germany). Images were acquired and processed in ZEISS ZEN software (Zen 3.5, Carl Zeiss Microscopy). Overview images were acquired using a 10x/0.45 objective, whereas high-resolution images were acquired using a 63x/1.4 oil-immersion objective. All images were collected using identical acquisition settings within each staining experiment to allow direct comparison between samples. Images were acquired at a resolution of 16 bits, and brightness and contrast were adjusted uniformly across all images.

#### Western blot

Specificity of the CA2 antibody in chicken tissue was validated by Western blotting of chicken retina and chicken kidney, with mouse kidney as a positive control (Spicer et al., 1979). Tissue samples (10-20 mg) were homogenised in RIPA buffer (50 mM Tris, 150 mM NaCl, 1% NP-40, 0.1% SDS) with complete miniprotease inhibitor cocktail (Cat# 04693159001, Roche, one tablet per 10 mL) using a Precellys homogeniser (2x30 s, 6500 rpm; Bertin Technologies, Montigny-le-Bretonneux, France) and protein concentration was determined using the Pierce™ 660 nm Protein Assay (Thermo Fisher Scientific). Samples were mixed 3:1 with 4x Laemmli sample buffer (Cat# 1610747, Bio-Rad) supplemented with dithiothreitol (DTT; final concentration 50 mM), denatured for 5 min at 95°C and separated on 10% Mini-PROTEAN TGX precast gels (Cat# 4561035, Bio-Rad, Hercules, CA, USA) in Tris/glycine/SDS running buffer (Cat# 1610732, Bio-Rad) at 70 V for 15 min followed by 130 V for 60 min. Proteins were transferred to 0.2 µm nitrocellulose membranes (Trans-Blot Turbo Midi, Cat# 1704159, Bio-Rad) using a Trans-Blot Turbo System (Bio-Rad) at 1.3 A and up to 25 V for 7 min. Membranes were blocked for 1 h at room temperature in blocking buffer (3% (w/v) BSA in TBST) and incubated overnight at 4°C with sheep anti-CA2 (see above) at 1 µg mL^−1^ (1:200) in blocking buffer. After washing in TBST (3 x 5 min), membranes were incubated for 1 h at room temperature in the dark, with donkey anti-sheep AF647 (see above) at 0.2 µg mL^−1^ (1:10,000), washed again, and imaged on an Odyssey XF imaging system (LI-COR Biosciences) using Image Studio software (v6.0, LI-COR Biosciences).

### Statistical analysis

Statistical analyses were performed in R (R Core Team, 2026). Plasma (or blood) and vitreous humour were compared using paired tests: normality of each group was assessed with Shapiro–Wilk tests, and a paired t-test was used when both groups were normally distributed (*P* > 0.05); otherwise, a Wilcoxon signed-rank test with normal approximation was used. β_NB_ was calculated for each sample as the slope of the linear regression of [HCO_3_^−^] against pH across the three pCO_2_ levels (eq. 2), and vitreous humour and plasma β_NB_ were compared using a paired t-test (n = 4 chickens). The relationships between β_NB_ and protein or phosphate concentration were assessed by linear regression for each individual, and the mean slope was compared with 0 using one-sample t-tests. Similarly, the slope of [HCO_3_^−^] against [Na^+^] or [Cl^−^] between plasma and vitreous humour was calculated for each individual, and the mean slope was compared with 1 using one-sample t-tests. Values in the text are given as mean and s.d. Differences were considered significant at *P* < 0.05.

## Results and discussion

### A compensated pH disturbance in the bird eye

*In vivo* measurements revealed differences in acid-base status between vitreous humour and plasma in chickens. The vitreous humour had a nearly threefold higher pCO_2_ (94 mmHg, s.d. = 14.9) than blood (33.8 mmHg, s.d. = 14.4; t_7_ = 6.9, n = 8, *P* < 0.001, paired t-test; Fig. 1E). Blood pCO_2_ values were within the expected physiological range for birds (Chiodi and Terman, 1965; Kawashiro and Scheid, 1975) and agree with previous vitreous and blood pCO_2_ values obtained by microsensor measurements and expiratory gas analysis, respectively (Damsgaard et al., 2026). The elevated vitreous pCO_2_ indicates that CO_2_ accumulates locally within the intraocular space, exceeding what can be explained by systemic blood pCO_2_ alone, reflecting high retinal CO_2_ production combined with limited vascular CO_2_ clearance.

**Fig. 1.**
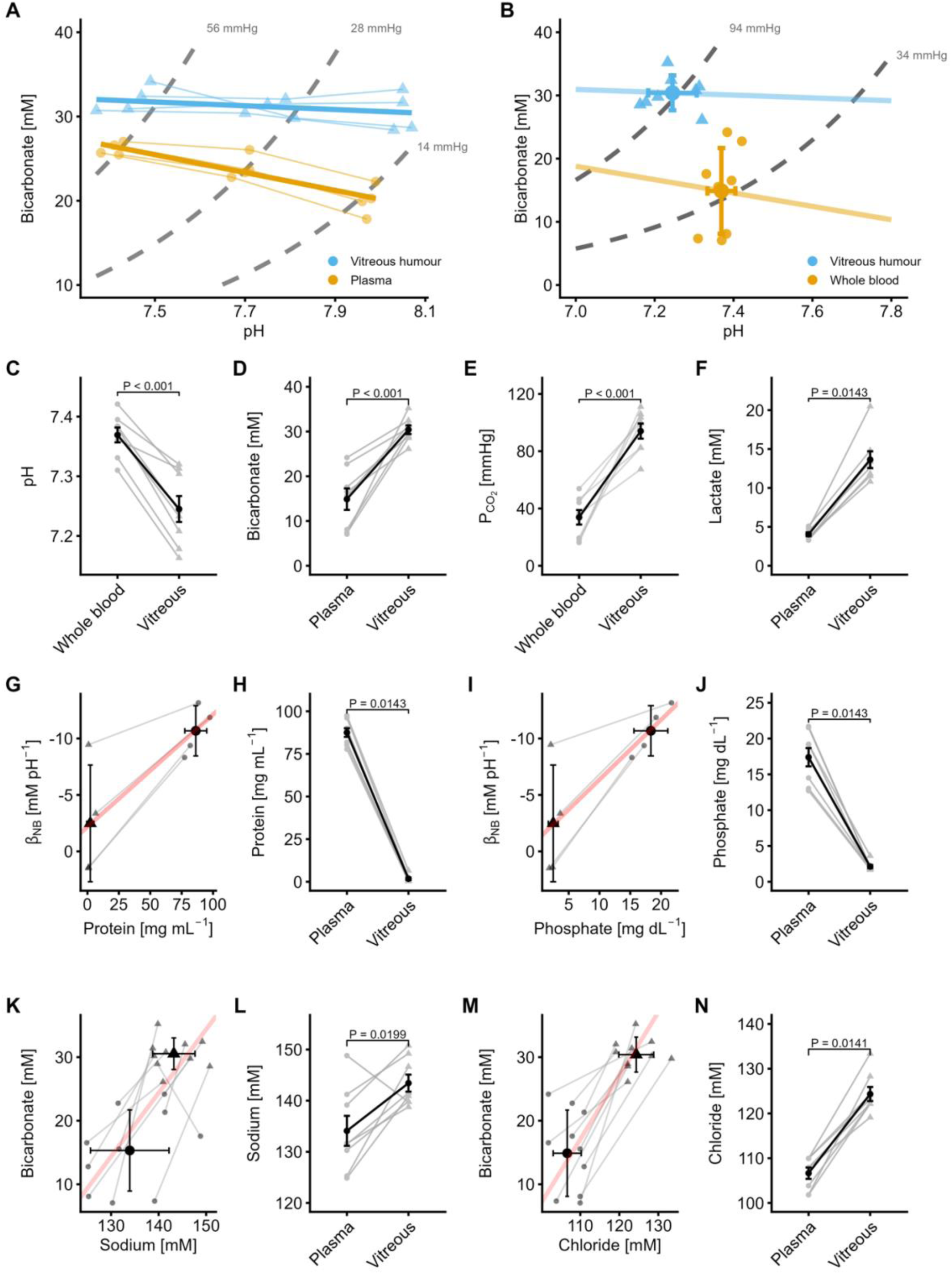
Acid-base balance and ionic composition in plasma and vitreous humour. **(A)** Davenport diagram showing buffer lines for vitreous (blue; n = 4) and plasma (yellow; n = 4), with pCO_2_ isopleths of 2% (14 mmHg), 4% (28 mmHg), and 8% (56 mmHg). Thin lines connect measurements from the same sample; thick lines show the linear regression across all samples within each group. **(B)** Davenport diagram from in vivo measurements (n = 8 hens; means ± s.d.), showing buffer lines for vitreous (blue) and blood (yellow) with slopes from the regression of the tonometry data for each group (A), with pCO_2_ isopleths at mean vitreous (94 mmHg) and mean blood (34 mmHg) values. **(C–F)** Measured pH, bicarbonate, pCO_2_, and lactate concentrations in blood and vitreous. **(G,I)** Relationship between protein or phosphate concentrations and β_NB_, with the red line showing the linear regression across all samples; note the reversed y-axis. **(H,J)** Measured concentrations of protein and phosphate in plasma and vitreous. **(K,M)** Correlation between sodium or chloride and bicarbonate concentrations, with the red line of slope = 1 indicating a 1:1 relationship. **(L,N)** Measured concentrations of sodium and chloride in plasma and vitreous.

Lactate concentration was similarly elevated in vitreous humour (13.6 mM, s.d. = 3.05) compared to plasma (4.0 mM, s.d. = 0.67; V = 36, n = 8, *P* = 0.014, Wilcoxon signed-rank test; Fig. 1F), consistent with anaerobic metabolism within the inner retina. Together, the elevated pCO_2_ and lactate accumulation represent a substantial combined acid load, yet vitreous pH (7.25, s.d. = 0.06) was only marginally lower than blood pH (7.37, s.d. = 0.04; t_7_ = −8.28, n = 8, *P* < 0.001, paired t-test; Fig. 1C). If blood had been exposed to similarly high pCO_2_ and lactate levels as found in the vitreous, pH would have been 6.8 (s.d. = 0.16, n = 5; eq. 5). In the remaining three hens, pH could not be calculated because the excess vitreous lactate exceeded blood [HCO_3_^−^]. Together, these results show that the bird eye largely regulates intraocular pH despite a sustained acid load.

To test whether non-bicarbonate buffers contribute markedly to vitreous pH buffering, we measured the *ex vivo* non-bicarbonate buffer capacity (β_NB_), i.e. the change in [HCO_3_^−^] per unit change in pH, which reflects buffering by proteins and phosphate. Vitreous humour had a fourfold lower β_NB_ (−2.65 mM per pH unit, s.d. = 5.17) than plasma (−10.8 mM per pH unit, s.d. = 2.27; t_3_ = 5.29, n = 4, *P* = 0.013, paired t-test; Fig. 1A,B).

The low vitreous β_NB_ appears to be linked to a low content of non-bicarbonate buffer groups, as both protein and phosphate levels were lower in the vitreous humour (protein: 1.77 mg mL^−1^, s.d. = 2.04; phosphate: 2.12 mg dL^−1^, s.d. = 0.62) compared to plasma (protein: 87.51 mg mL^−1^, s.d. = 7.17; V = 0, n = 8, *P* = 0.014; phosphate: 17.39 mg dL^−1^, s.d. = 3.61; V = 0, n = 8, *P* = 0.014, Wilcoxon signed-rank tests; Fig. 1H,J). We quantified the relationship between β_NB_ and protein or phosphate concentration across both vitreous and plasma samples from the tonometer experiments. This demonstrated a consistent association between the concentrations of both non-bicarbonate buffers and the magnitude of β_NB_ (protein: mean slope = −0.099, t_3_ = −4.75, *P* = 0.018; phosphate: mean slope = −0.549, t_3_ = −4.29, *P* = 0.023, one-sample *t*-tests; Fig. 1G,I). Thus, the low β_NB_ of vitreous humour, explained by its poor protein and phosphate content, suggests that the eye does not rely on non-bicarbonate buffers for pH defence but on active transepithelial transport of acid-base-relevant ions.

[Na^+^] was higher in the vitreous (143 mM, s.d. = 4.71) than in plasma (134 mM, s.d. = 8.30; t_7_ = 3.0, n = 8, *P =* 0.019, paired *t*-test; Fig. 1L), and [Cl^−^] was similarly elevated in vitreous (124 mM, s.d. = 4.46) compared to plasma (106 mM, s.d. = 3.59; V = 36, n = 8, *P* = 0.014, Wilcoxon signed-rank test; Fig. 1N). [HCO_3_^−^] was higher in vitreous (30 mM, s.d. = 2.73) compared to the plasma (15 mM, s.d. = 6.75; t_7_ = 5.8, n = 8, *P* < 0.001, paired *t*-test; Fig. 1D), directly supporting HCO ^−^ as the primary buffer in the vitreous. To further investigate potential transport of HCO_3_^−^ into the vitreous, the proportionality between HCO_3_^−^ and Na^+^ or Cl^−^ was examined by individual linear regressions. One-sample *t*-tests confirmed that the mean slopes did not deviate significantly from 1 for either Cl^−^/HCO_3_^−^ (mean slope: 0.94; t_7_ = −0.30, *P* = 0.77; Fig. 1M) or Na^+^/HCO_3_^−^ (mean slope: 1.15; t_7_ = 0.33, *P* = 0.74; Fig. 1K). On average, HCO_3_^−^ thus maintains a proportional relationship near 1:1 with both Na^+^ and Cl^−^, indicating that Na^+^ and Cl^−^ act as acid-base-relevant counterions. The close co-variation between Na^+^ and HCO_3_^−^ is consistent with the activity of Na^+^/HCO_3_^−^ cotransporters of the SLC4 family, which are well-established mediators of active HCO_3_^−^ transport across cell membranes (Boron, 2004; Holmberg et al., 2024), and have been described in ocular tissues of humans, rats and mice (Bok et al., 2001; Seki et al., 2013).

A comparable mechanism, potentially localised to the pecten oculi, could explain the vitreous HCO_3_^−^ accumulation. It is also possible that the elevated Na⁺ concentration is unrelated to HCO_3_^−^ and instead reflects activity of a Na^+^/H^+^ exchanger, such as those from the SLC9 family, which may contribute to pH regulation by extruding H^+^ from the vitreous (Xu et al., 2018). The parallel relationship between Cl^−^ and HCO_3_^−^ is unexpected, as members of the SLC4 and SLC26 transporter families typically function as Cl^−^/HCO_3_^−^ antiporters, predicting lower Cl^−^ where HCO_3_^−^ is elevated (Boron, 2004). The observed co-directional pattern may instead reflect coupled transport, although simultaneous inward flux would require compensation by other ions to maintain charge balance. Together, the low vitreous β_NB_, the elevated vitreous HCO_3_^−^ concentration and its near 1:1 co-variation with Na^+^ indicate that the bird eye relies on active bicarbonate transport, providing a continuous influx of HCO_3_^−^ for intraocular pH defence.

### The pecten regulates vitreous pH

To identify acid-base-relevant transporters and enzymes in the blood-retinal barrier, we used a spatial transcriptomics dataset of the chicken eye. Using this dataset, we identified CA2, CA4, and two V-ATPase subunits (ATP6V1C2 and ATP6V0E1) as the primary pH-regulatory genes enriched in the pecten (Fig. 2A–D). CA4 and both V-ATPase subunits showed strong pecten-specific upregulation (CA4: log_2_FC = 5.84, −log(p) = 109, n = 1; Fig. 2B; ATP6V1C2: log_2_FC = 4.69, −log(p) = 140, n = 1; Fig. 2C; ATP6V0E1: log_2_FC = 2.437, −log(p) = 51, n = 1; Fig. 2D), while CA2, despite its broader ocular expression (log_2_FC = 1.24, −log(p) = 21, n = 1; Fig. 2A), was included due to its consistently high expression and established role in intracellular pH regulation.

**Fig. 2.**
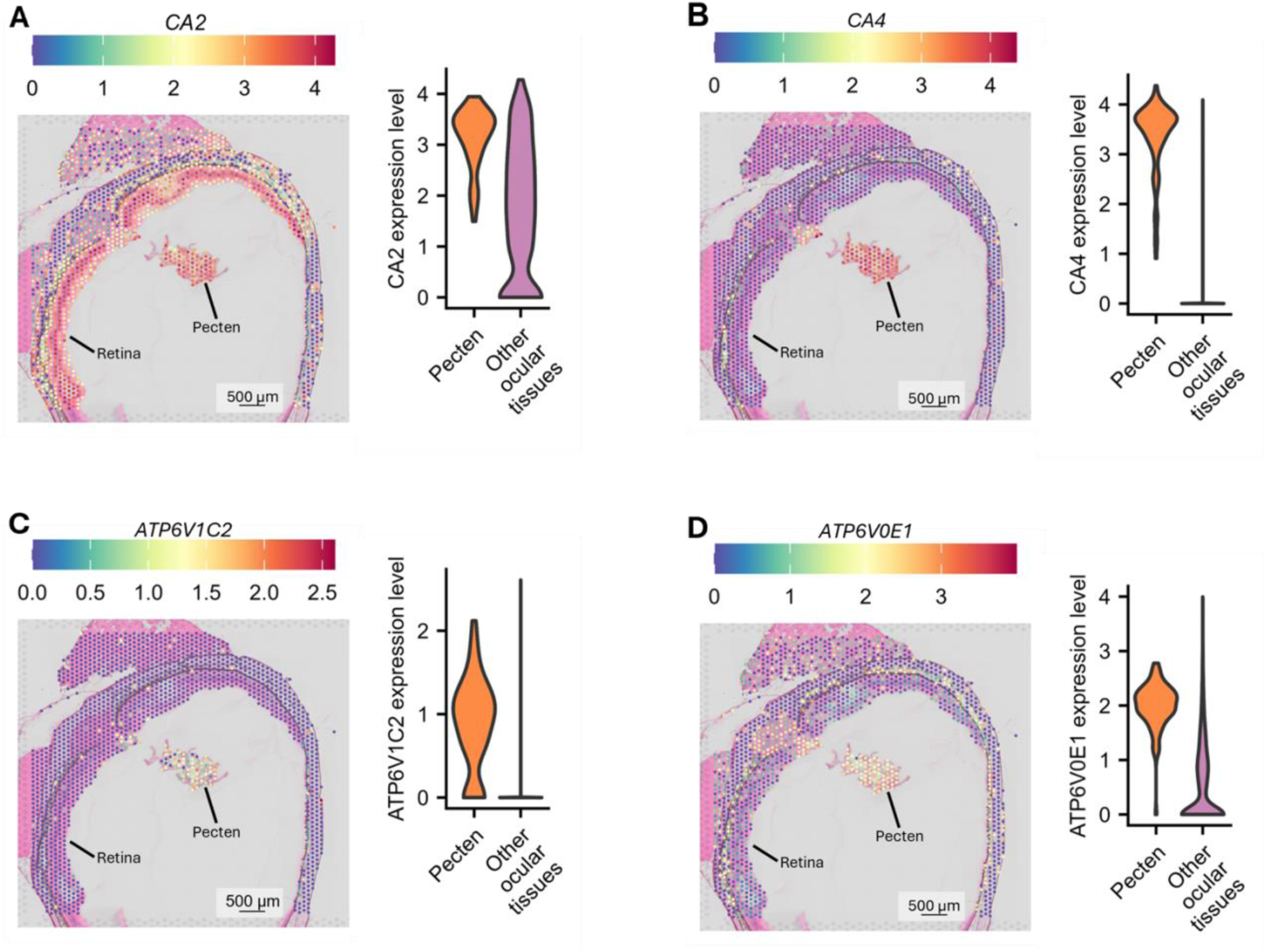
Spatial expression of acid-base regulatory genes in the chicken eye. **(A–D)** Spatial expression maps of CA2, CA4, ATP6V1C2, and ATP6V0E1 across the eye section. Each spot represents a transcript-capturing location on the Visium slide and is colour-coded according to local gene expression intensity. For each gene, the corresponding violin plot on the right compares expression levels in the pecten oculi (orange) with other ocular tissues (purple). The width of each violin reflects the density of expression values, with wider regions indicating a higher concentration of data points.

Notably, no members of the classic Na^+^/HCO_3_^−^, Na^+^/H^+^, or Cl^−^/HCO_3_^−^ transporter families (SLC4, SLC9, and SLC26) showed significant enrichment in the pecten. This was unexpected given the elevated vitreous Na^+^, Cl^−^, and HCO_3_^−^ concentrations reported above, which would otherwise suggest active ion transport. Among other ocular tissues, only SLC4A10, encoding a sodium-coupled bicarbonate transporter, showed notable enrichment, consistent with its established role in neuronal pH regulation (Hilgen et al., 2012). SLC9A3 showed a high log-fold change in the pecten but was detected in only 11.5% of pecten spots (Table S1). This suggests that SLC9A3 and its regulatory cofactors (SLC9A3R1 and SLC9A3R2) are either of limited functional relevance in the pecten or confined to a small subset of pecten cell types (Figs S2–S6).

Immunohistochemistry confirmed CA2 protein in the pecten, validating that the transcriptomic signal detected at the mRNA level translates into protein expression. CA2 immunoreactivity was most prominent at the periphery of the pecten (Fig. 3A; Fig. S7), consistent with its known cytosolic localisation in vascular endothelial cells (McMurtrie et al., 2004). However, because bird red blood cells are nucleated, intravascular blood cells cannot be distinguished from pecten tissue by DAPI staining alone (Nikinmaa, 2012). Part of the immunoreactivity may therefore reflect CA2 in circulating red blood cells rather than in the pecten endothelium. Co-staining with endothelial-specific markers in future studies would help to resolve this. Whereas specificity of the CA2 antibody in chicken tissue was confirmed by Western blotting (Fig. S1), the available antibodies against CA4 and V-ATPase could not be validated; their presence in the pecten is therefore based on mRNA expression (Fig. 2B–D).

**Fig. 3.**
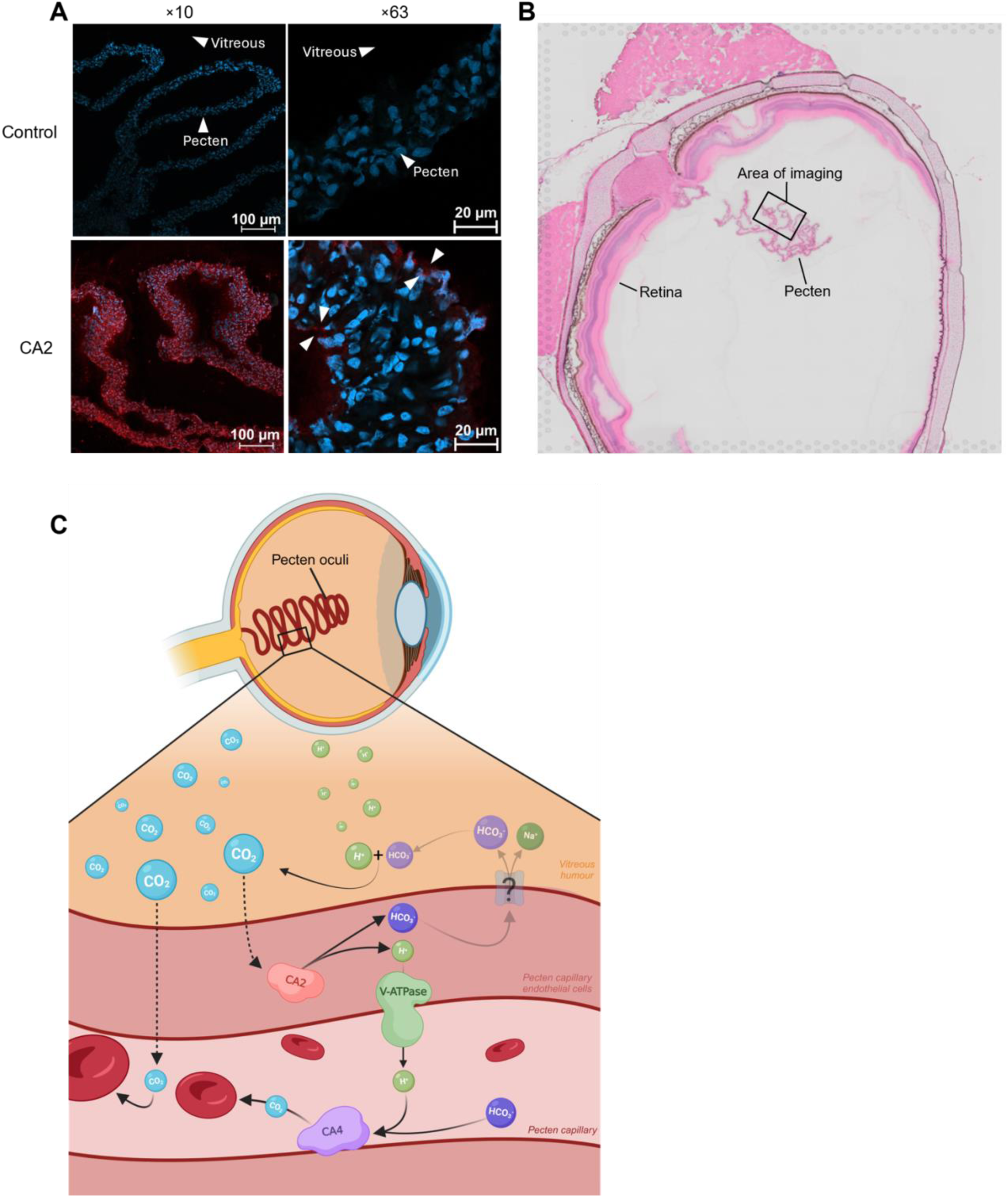
Protein localisation and proposed CO_2_ transport model in the bird pecten oculi. **(A)** Immunofluorescence images of the chicken (*Gallus gallus domesticus*) pecten oculi at x10 (left) and x63 (right) magnification. Top row: control (DAPI, blue). Bottom row: CA2 (red) merged with DAPI, showing CA2 present at the periphery of the pecten, consistent with localisation in endothelial cells (arrowheads) (n = 3). **(B)** Raw tissue section indicating the imaged area. **(C)** Proposed model for CO_2_ removal from the vitreous humour via the pecten oculi. Dashed arrows indicate diffusion, solid arrows reactions or transport, and “?” an unidentified transporter. The positions of CA4 and V-ATPase are based on spatial transcriptomics (mRNA) data only. The inset (top right) illustrates the intraretinal O_2_ gradient, with CO_2_ produced by the oxygenated outer retina and lactate by the anoxic inner retina.

Based on the combined gene expression data and CA2 protein localisation, we suggest a putative mechanism for pecten pH regulation (Fig. 3C). Two mechanisms may account for CO_2_ removal via the pecten. First, CO_2_ may simply diffuse across the full length of the pecten and be taken up directly by the capillary blood. Alternatively, a faster route may exist at the periphery, where CA2-expressing endothelial cells may facilitate rapid intracellular conversion of diffusible CO_2_ into HCO_3_^−^ and H^+^, thereby maintaining the CO_2_ diffusion gradient from the vitreous into the pecten (Fig. 3C). The resulting H^+^ could subsequently be extruded by V-ATPase from the endothelial cells into the capillary lumen. Here, membrane-bound CA4 may catalyse the interconversion of HCO_3_^−^ and CO_2_ in the plasma, allowing CO_2_ uptake by circulating red blood cells (Fig. 3C). In parallel, HCO ^−^ generated by CA2 may be transported toward the vitreous, contributing to the HCO_3_^−^ buffering capacity observed in this compartment. Such HCO_3_^−^ transport would classically involve Na^+^/HCO_3_^−^ cotransporters, a mechanism that would be consistent with the elevated Na^+^ concentrations detected in the vitreous. However, the transporter responsible for this HCO_3_^−^ flux remains to be identified, as none of the classical SLC4 or SLC26 family members were enriched in the pecten.

## Conclusion

Our data demonstrate a substantial intraocular acid load arising from elevated pCO_2_ and lactic acid, buffered by bicarbonate. We further identify the pecten oculi as a key blood-retinal barrier facilitating the necessary exchange of acid-base equivalents to regulate intraocular pH. Together with the previous identification of the pecten as a metabolite exchange epithelium (Damsgaard et al., 2026), these data position the pecten oculi as a hub for metabolite exchange and pH regulation, connecting the blood and the retina via the vitreous humour. These findings point to the pecten as a key evolutionary adaptation enabling the uniquely low vascularisation and high visual performance of the bird retina. Future studies involving functional inhibition of CA2 and CA4, targeted investigation of bicarbonate transporters, and antibodies validated for avian CA4 and V-ATPase will be essential to fully resolve the mechanisms underlying intraocular acid-base regulation.

## Acknowledgements

We thank Kathrin Hinz, Emily Leishmann, Heidi M. Jensen, Rasmus D. Buchanan, Per G. Henriksen, Simon E. Olsen, and Julius Gaarden for assistance with animal handling and sampling, Helene M. Andersen and Trine W. Mikkelsen for valuable advice on cryosectioning, and Elin E. Petersen for laboratory support. The authors acknowledge the Bioimaging Core Facility, Health, Aarhus University, Denmark, for the use of equipment and support.

Fig. 3C was created in BioRender. Mikkelsen, A. V. (2027) https://BioRender.com/vr29tax.

## Author contributions

Conceptualisation: A.V.G.T.M., T.V-J., T.W., C.J.A.W., C.D.

Formal analysis: A.V.G.T.M., F.T.K.

Data curation: A.V.G.T.M.

Funding acquisition: C.D., C.J.A.W.

Methodology: A.V.G.T.M., M.H.F., A.B., F.T.K., C.J.A.W.

Project administration: C.D.

Resources: T.V-J., T.W., C.J.A.W., C.D.

Supervision: M.H.F., T.W., C.D.

Writing – original draft: A.V.G.T.M.

Writing – review & editing: All authors

## Data availability

Data will be deposited on GitHub at submission.

## Supplementary material

**Fig. S1.**
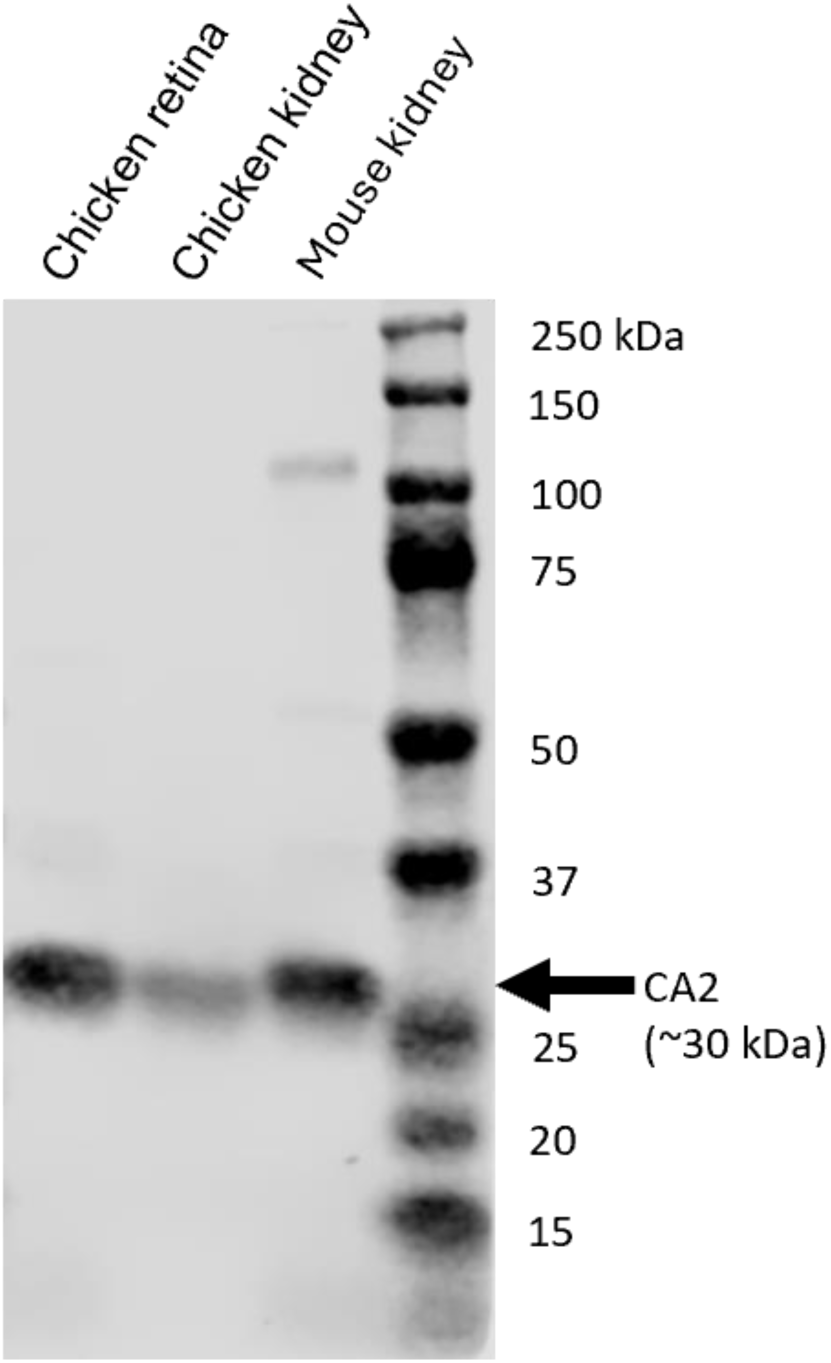
Western blot validation of the anti-CA2 antibody in chicken tissue. Western blot of chicken retina, chicken kidney and mouse kidney (positive control) probed with sheep anti-CA2. A single band at the expected molecular weight of CA2 (∼30 kDa; arrow) was detected in all samples. Molecular weight markers (kDa) are indicated on the right.

**Fig. S2.**
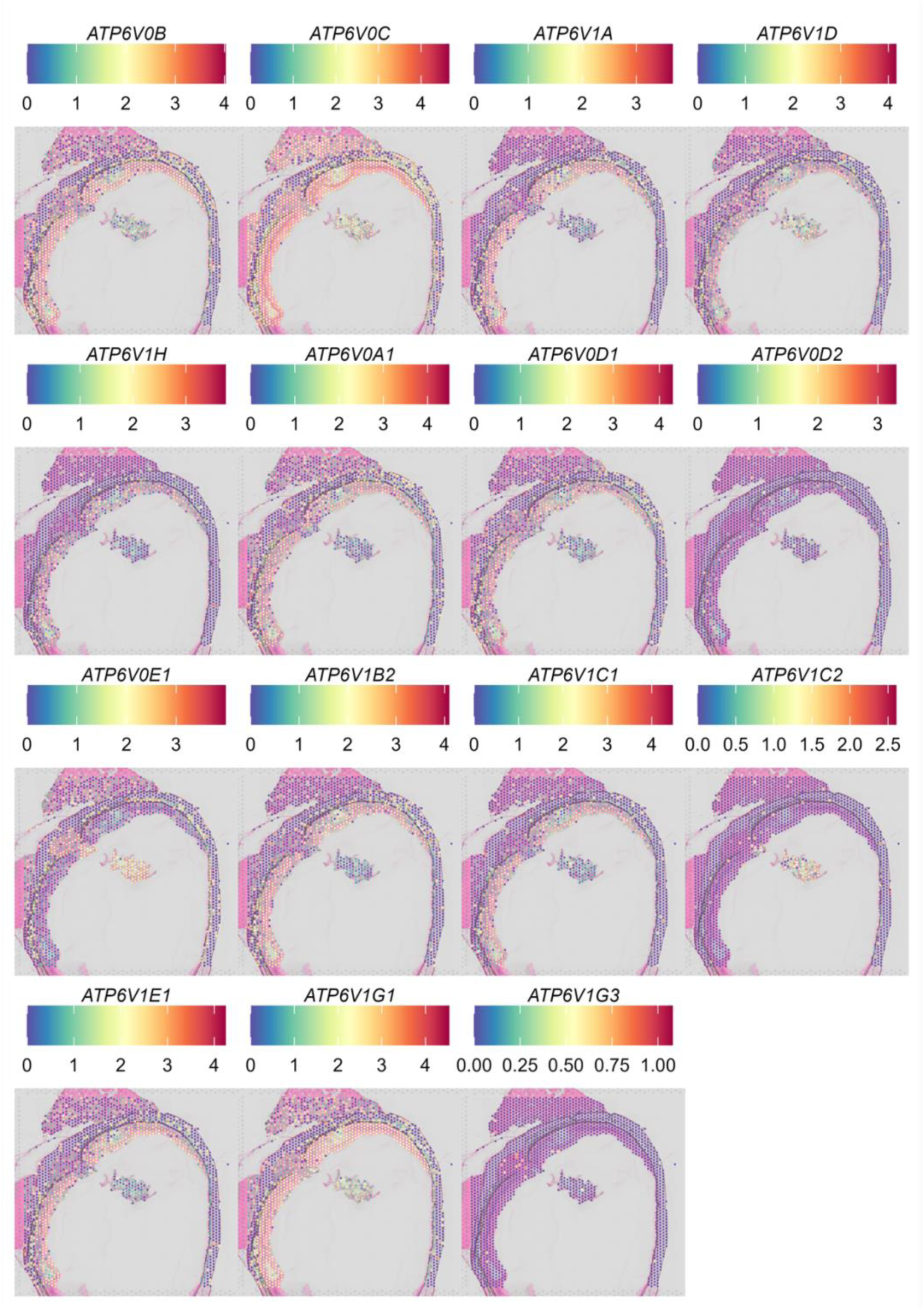
Spatial expression of Vacuolar-type H^+^-ATPase (V-ATPase) in chicken eye. Expression levels of the investigated V-ATPases mapped onto cryosections of chicken eye. Each spot represents a gene expression signal and is colour-coded according to expression intensity.

**Fig. S3.**
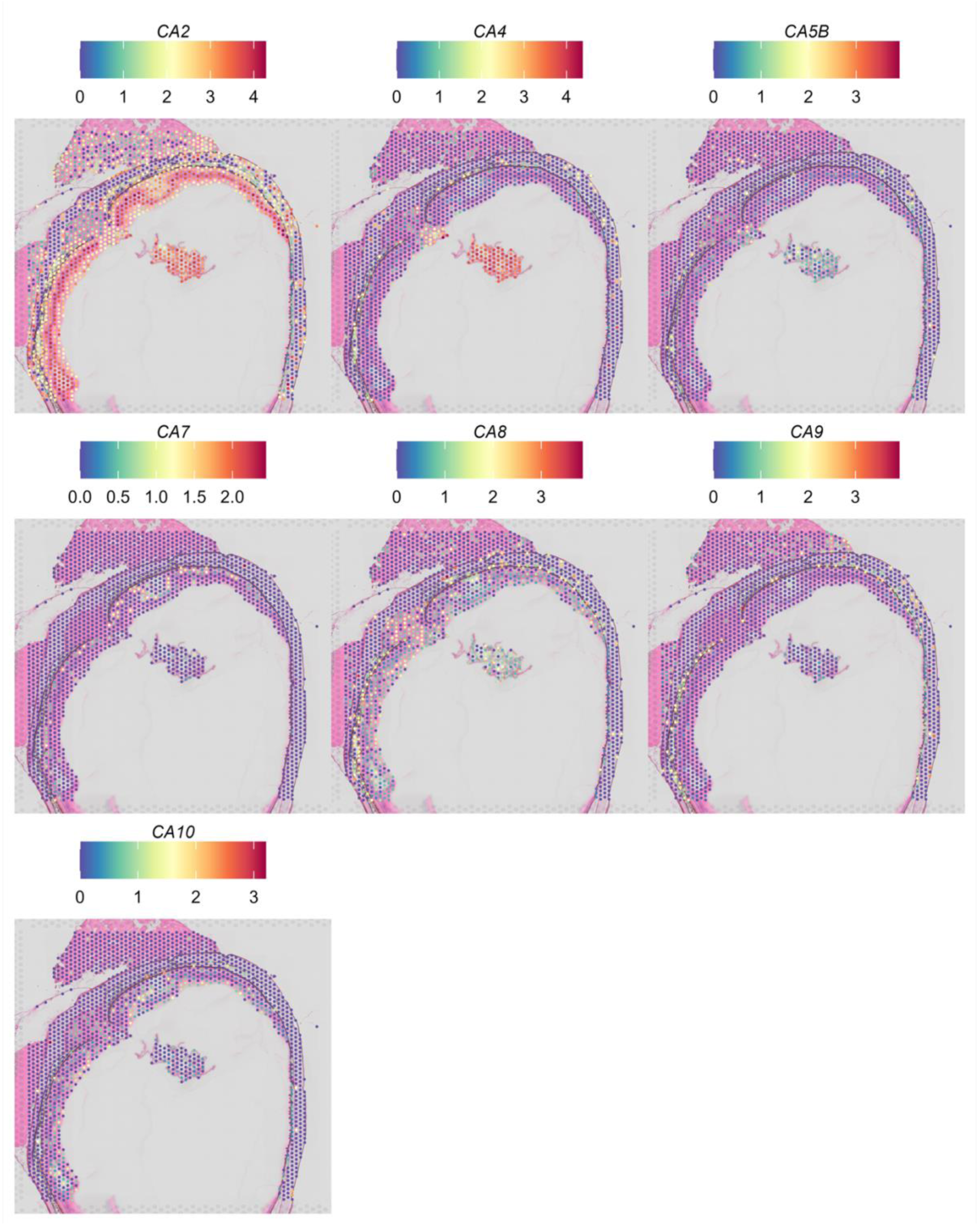
Spatial expression of Carbonic Anhydrase (CA) isoforms in chicken eye. Expression levels of the investigated CA isoforms mapped onto cryosections of chicken eye. Each spot represents a gene expression signal and is colour-coded according to expression intensity.

**Fig. S4.**
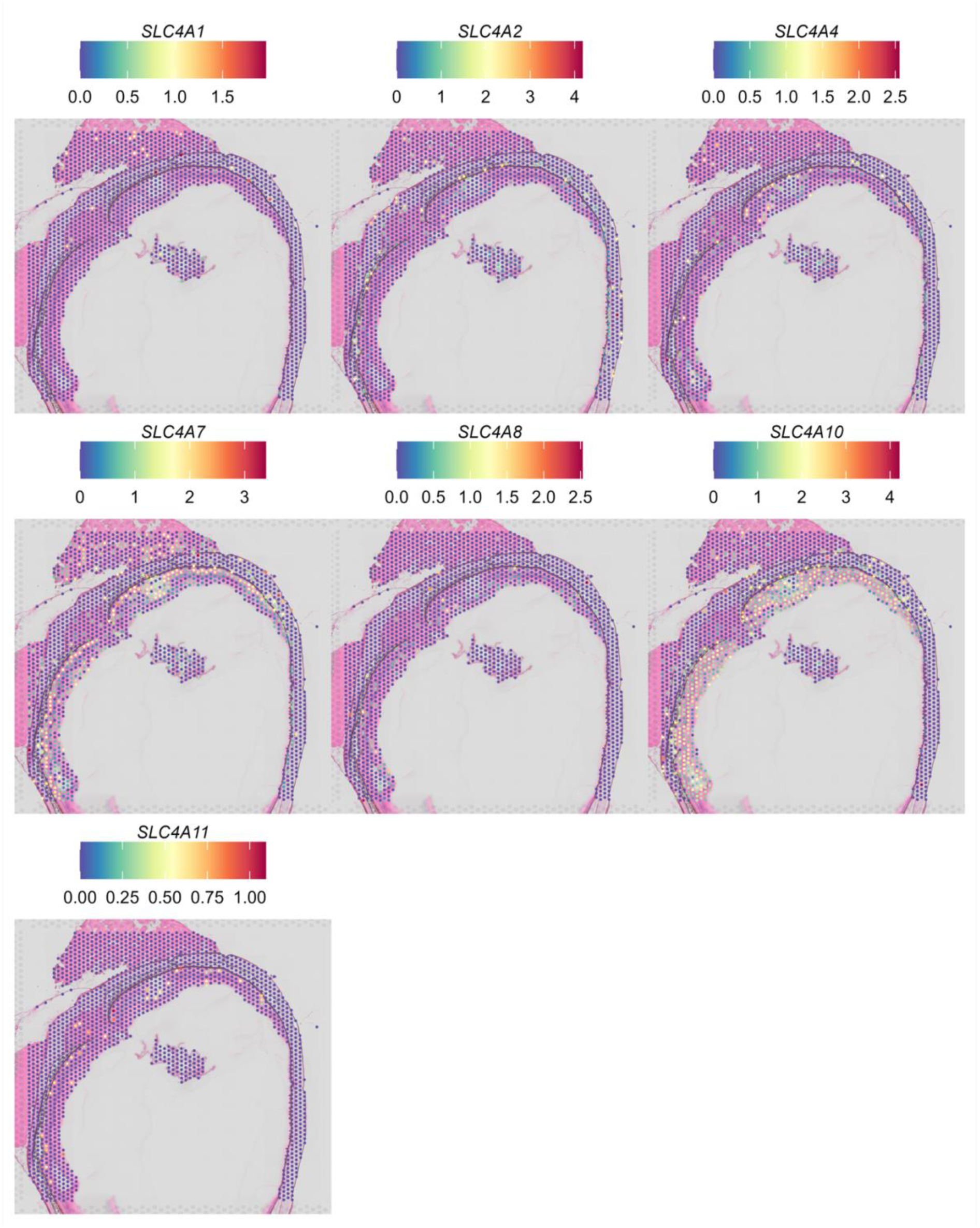
Spatial expression of Solute Carrier Family 4 (SLC4) in chicken eye. Expression levels of the investigated SLC4 mapped onto cryosections of chicken eye. Each spot represents a gene expression signal and is colour-coded according to expression intensity.

**Fig. S5.**
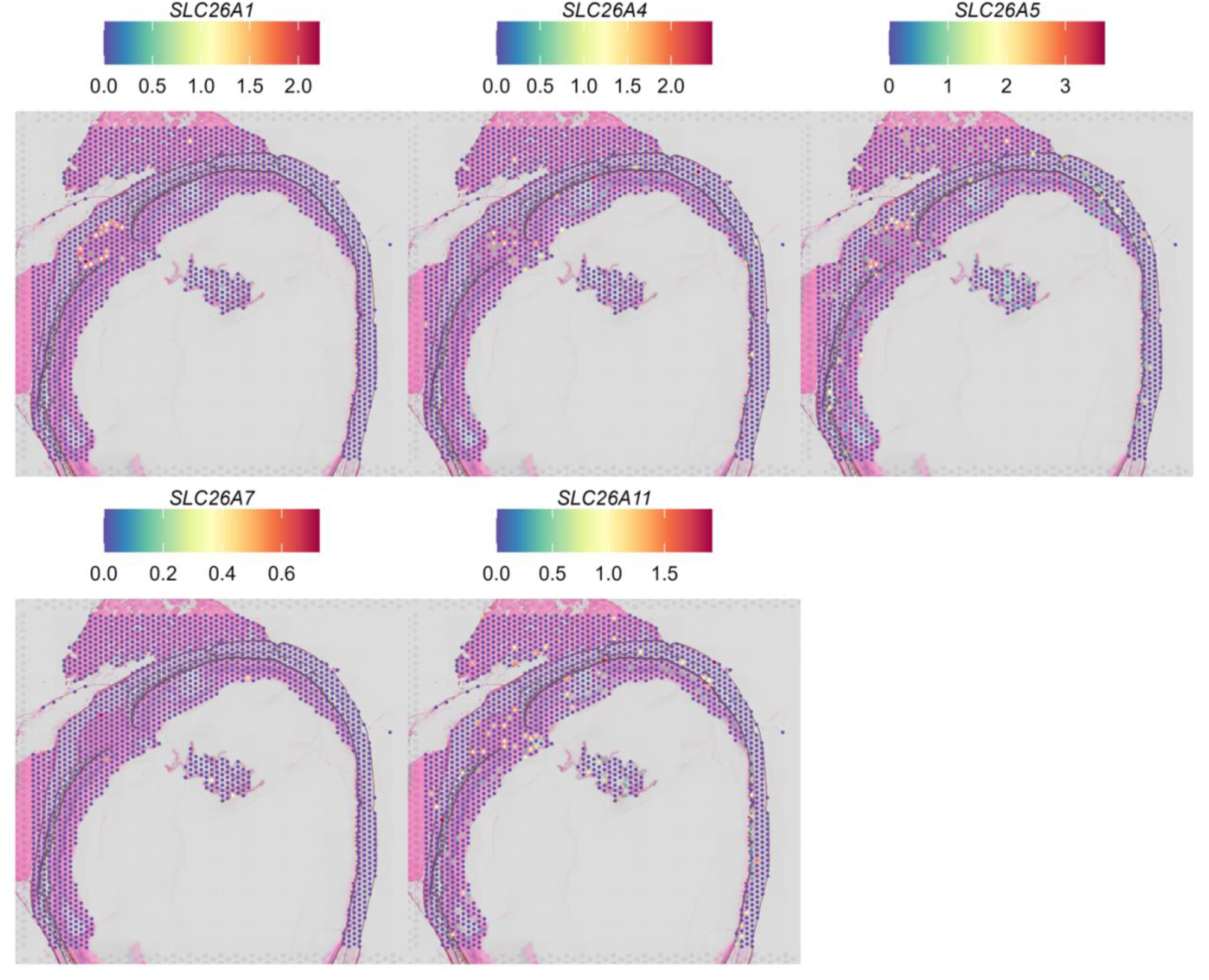
Spatial expression of Solute Carrier Family 26 (SLC26) in chicken eye. Expression levels of the investigated SLC26 mapped onto cryosections of chicken eye. Each spot represents a gene expression signal and is colour-coded according to expression intensity.

**Fig. S6.**
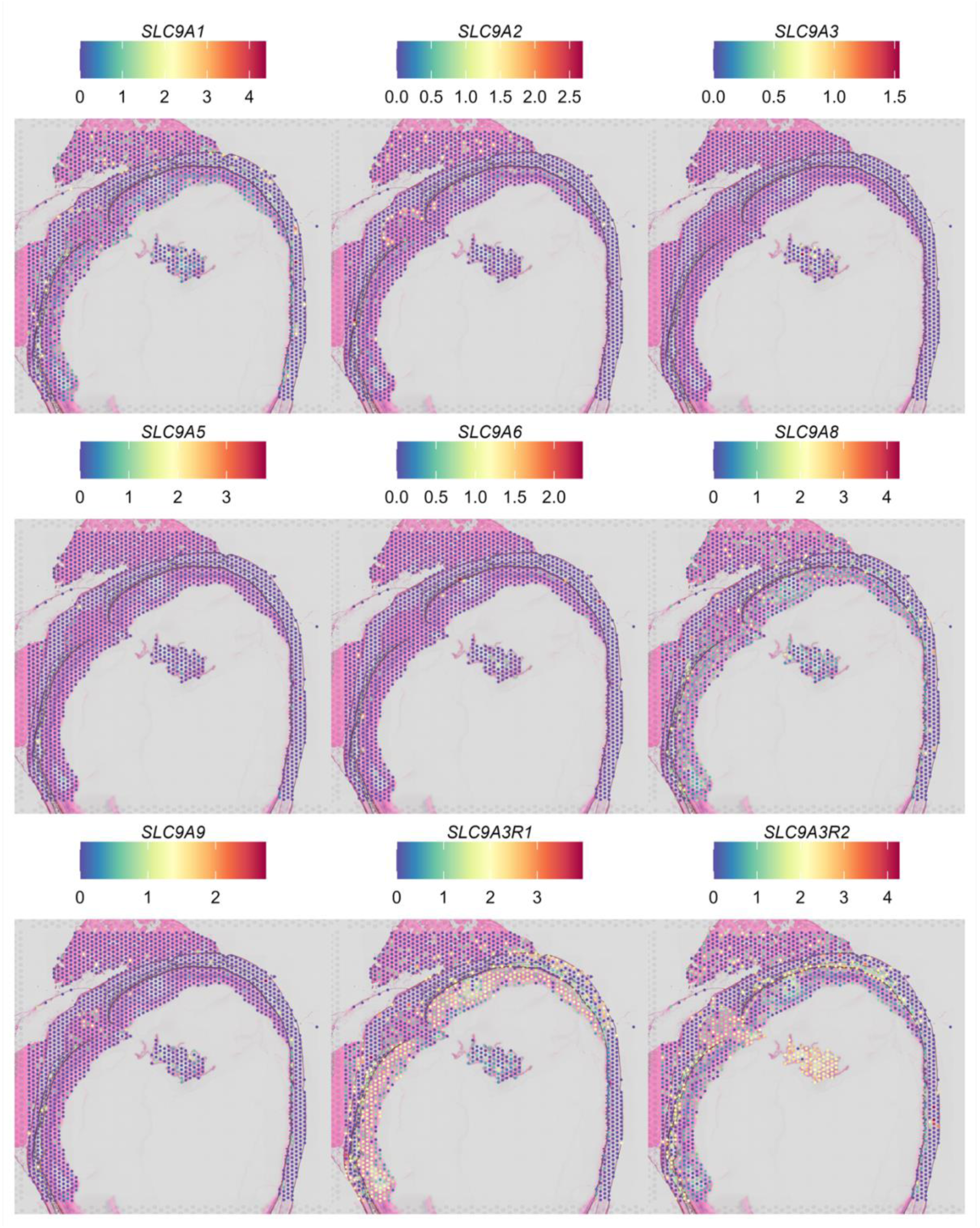
Spatial expression of Solute Carrier Family 9 (SLC9) in chicken eye. Expression levels of the investigated SLC9 isoforms mapped onto cryosections of chicken eye. Each spot represents a gene expression signal and is colour-coded according to expression intensity.

**Fig. S7.**
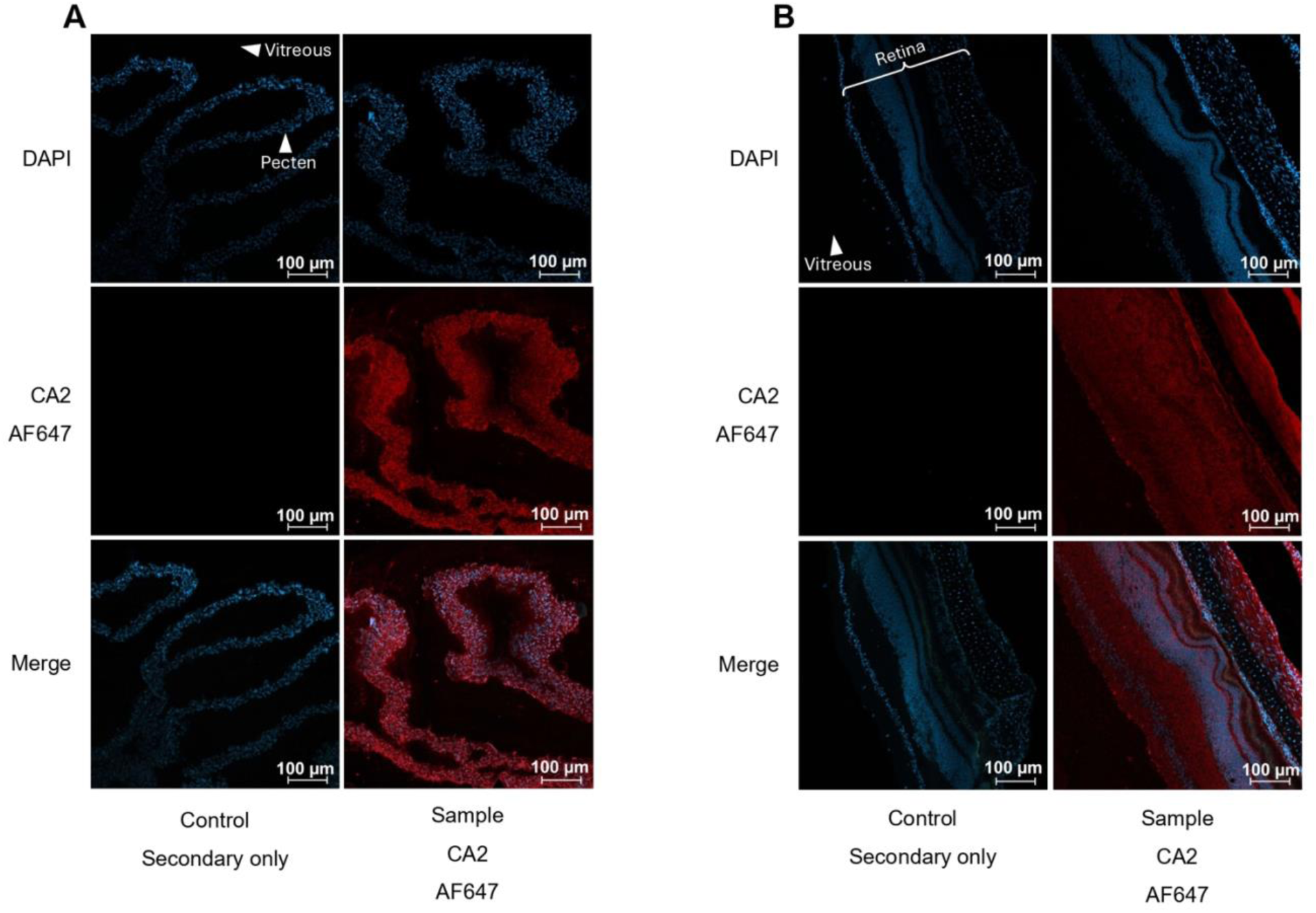
Immunofluorescence localisation of CA2 in the chicken pecten oculi and retina (10x magnification). Representative sections of **(A)** pecten oculi and **(B)** retina. For each tissue, the left column shows a secondary antibody-only control (secondary only) and the right column shows staining with anti-CA2 (anti-CA2). Rows show DAPI (blue; nuclei), CA2 (red; AF647) and the merged image.

**Table S1.** All genes investigated in relation to pH regulation in the pecten. Genes investigated in the pecten for roles in pH regulation. NA indicates genes not detected in the tissue; no images are available for these.

| **V-ATPase** |  |  |  |  |
|---|---|---|---|---|
| ***Gene*** | ***Log<sub>2</sub>FC*** | ***Pct<sub>pecten</sub>*** | ***Pct<sub>eye</sub>*** | ***−log(p)*** |
|---|---|---|---|---|
| **ATP6V0B** | −0.676 | 0.854 | 0.637 | 0 |
| **ATP6V0C** | −0.885 | 0.969 | 0.795 | 0 |
| **ATP6V1A** | −1.006 | 0.583 | 0.453 | 0 |
| **ATP6V1D** | 0.790 | 0.875 | 0.494 | 10 |
| **ATP6V1F** | *NA* | *NA* | *NA* | *NA* |
| **ATP6V1H** | −0.840 | 0.396 | 0.325 | 0 |
| **ATP6V0A1** | −1.440 | 0.646 | 0.590 | 0 |
| **ATP6V0A2** | *NA* | *NA* | *NA* | *NA* |
| **ATP6V0D1** | −0.827 | 0.708 | 0.522 | 0 |
| **ATP6V0D2** | −0.231 | 0.125 | 0.114 | 0 |
| **ATP6V0E1** | 2.437 | 0.990 | 0.468 | 51 |
| **ATP6V0E2** | *NA* | *NA* | *NA* | *NA* |
| **ATP6V1B2** | −1.319 | 0.688 | 0.501 | 0 |
| **ATP6V1C1** | −0.739 | 0.615 | 0.420 | 0 |
| **ATP6V1C2** | 4.686 | 0.823 | 0.046 | 140 |
| **ATP6V1E1** | −0.811 | 0.698 | 0.533 | 0 |
| **ATP6V1G1** | −0.140 | 0.990 | 0.658 | 0 |
| **ATP6V1G3** | 0.700 | 0.010 | 0.011 | 0 |
| **Carbonic anhydrase (CA) isoforms** |  |  |  |  |
| ***Gene*** | ***Log<sub>2</sub>FC*** | ***Pct<sub>pecten</sub>*** | ***Pct<sub>eye</sub>*** | ***−log(p)*** |
| **CA2** | 1.242 | 1.000 | 0.760 | 21 |
| **CA4** | 5.837 | 1.000 | 0.160 | 109 |
| **CA5B** | 2.224 | 0.719 | 0.129 | 49 |
| **CA7** | −0.418 | 0.062 | 0.062 | 0 |
| **CA8** | 0.717 | 0.781 | 0.363 | 11 |
| **CA9** | −2.564 | 0.104 | 0.157 | 0 |
| **CA10** | −0.291 | 0.229 | 0.180 | 0 |
| **CA13** | *NA* | *NA* | *NA* | *NA* |
| **CA14** | *NA* | *NA* | *NA* | *NA* |
| **SLC4 family** |  |  |  |  |
| ***Gene*** | ***Log<sub>2</sub>FC*** | ***Pct<sub>pecten</sub>*** | ***Pct<sub>eye</sub>*** | ***−log(p)*** |
| **SLC4A1** | 0.581 | 0.052 | 0.017 | 0 |
| **SLC4A2** | −1.060 | 0.146 | 0.095 | 0 |
| **SLC4A4** | −0.766 | 0.062 | 0.076 | 0 |
| **SLC4A7** | −1.974 | 0.135 | 0.297 | 0 |
| **SLC4A8** | −2.735 | 0.000 | 0.062 | 0 |
| **SLC4A10** | −3.275 | 0.083 | 0.362 | 4 |
| **SLC4A11** | −1.012 | 0.000 | 0.025 | 0 |
| **SLC26 family** |  |  |  |  |
| ***Gene*** | ***Log<sub>2</sub>FC*** | ***Pct<sub>pecten</sub>*** | ***Pct<sub>eye</sub>*** | ***−log(p)*** |
| **SLC26A1** | −1.096 | 0.010 | 0.016 | 0 |
| **SLC26A2** | *NA* | *NA* | *NA* | *NA* |
| **SLC26A4** | 0.134 | 0.031 | 0.036 | 0 |
| **SLC26A5** | −0.396 | 0.198 | 0.116 | 0 |
| **SLC26A6** | *NA* | *NA* | *NA* | *NA* |
| **SLC26A7** | 3.115 | 0.021 | 0.002 | 0 |
| **SLC26A9** | *NA* | *NA* | *NA* | *NA* |
| **SLC26A11** | 1.338 | 0.219 | 0.067 | 3 |
| **SLC9 family** |  |  |  |  |
| ***Gene*** | ***Log<sub>2</sub>FC*** | ***Pct<sub>pecten</sub>*** | ***Pct<sub>eye</sub>*** | ***−log(p)*** |
| **SLC9A1** | −0.438 | 0.354 | 0.249 | 0 |
| **SLC9A2** | −2.551 | 0.010 | 0.065 | 0 |
| **SLC9A3** | 7.434 | 0.115 | 0.000 | 33 |
| **SLC9A5** | −0.659 | 0.031 | 0.015 | 0 |
| **SLC9A6** | 1.199 | 0.062 | 0.013 | 0 |
| **SLC9A7** | *NA* | *NA* | *NA* | *NA* |
| **SLC9A8** | −0.349 | 0.417 | 0.351 | 0 |
| **SLC9A9** | 1.268 | 0.167 | 0.053 | 1 |
| **SLC9A3R1** | −1.875 | 0.438 | 0.465 | 0 |
| **SLC9A3R2** | 2.727 | 0.979 | 0.435 | 51 |

